# Parallel functional subnetworks embedded in the macaque face patch system

**DOI:** 10.1101/2021.10.25.465775

**Authors:** Soo Hyun Park, Kenji W. Koyano, Brian E. Russ, Elena N. Waidmann, David B. T. McMahon, David A. Leopold

**Author notes:** Center for Biomedical Imaging and Neuromodulation, Nathan Kline Institute; Orangeburg, USA. Department of Neuroscience, Icahn School of Medicine at Mount Sinai; New York City, USA. Department of Psychiatry, New York University at Langone; New York City, USA. Laboratory of Neurogenetics of Language, The Rockefeller University; New York City, USA.

## Abstract

During normal vision, our eyes provide the brain with a continuous stream of useful information about the world. How visually specialized areas of the cortex, such as face-selective patches, operate under natural modes of behavior is poorly understood. Here we report that, during the free viewing of videos, cohorts of face-selective neurons in the macaque cortex fractionate into distributed and parallel subnetworks that carry distinct information. We classified neurons into functional groups based on their video-driven coupling with fMRI time courses across the brain. Neurons from each group were distributed across multiple face patches but intermixed locally with other groups at each recording site. These findings challenge prevailing views about functional segregation in the cortex and underscore the importance of naturalistic paradigms for cognitive neuroscience.

**One-Sentence Summary:** Natural visual experience reveals parallel functional subnetworks of neurons embedded within the macaque face patch system

## Main Text

Face patches in the temporal cortex of macaques are spatially separate, densely interconnected (*1, 2*) regions, defined by their shared preference for faces (*3, 4*). Similar regions exist in the cortex of humans (*5*) and marmosets (*6*), suggesting this functional network is a conserved feature in primates (*7–9*). Individual patches differ in their neural response selectivity to facial attributes, such as viewpoint, identity, movement and expression (*10–12*). These differences have prompted speculation that face patches are arranged in one or more functional hierarchies (*13–15*).

Knowledge about the organization of the macaque face network derives principally from neural responses to briefly presented images. However, a few studies have begun to characterize the brain’s responses to freely viewed natural videos (*16–20*), with some evidence suggesting that the basic layout of the face patch network is preserved under these conditions (*17*). While naturalistic paradigms sacrifice precise control over visual stimulation, they are able to more closely approximate the conditions under which the visual specialization evolved in the primate brain (*21*). An important question is whether the principles derived from image responses will extend to more natural modes of vision (*22*).

Here we approach this question using a combination of methods that offers a new perspective on the visual responses of face-selective neurons. Briefly, we describe the activity of isolated single neurons in multiple nodes of the macaque face patch network during the free and repeated viewing of videos depicting natural social behaviors. Rather than taking the conventional approach of analyzing a neuron’s representation of visual features, we instead compare each neuron’s activity to corresponding fMRI responses across the brain obtained to the same video. (**Fig. 1A**). Thus, each neuron is characterized through its cortical network profile, summarized as a map of positive and negative covariations, rather than through its explicit stimulus selectivity. This approach provides a robust, data-driven method to assess the functional organization of the face patch system amid the complexity of naturalistic stimuli, in the context of the whole-brain network.

**Fig. 1.**
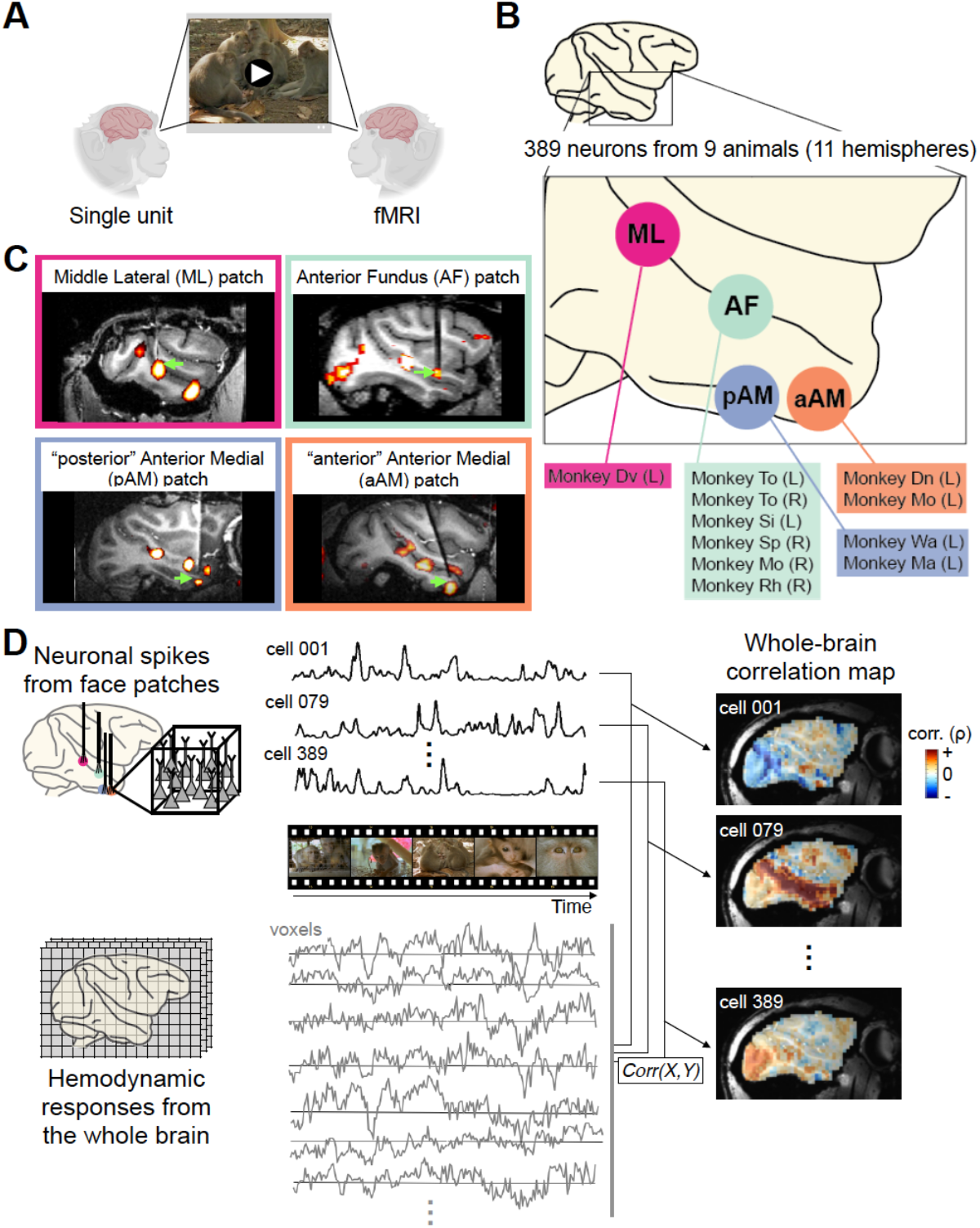
Computing whole-brain correlation maps for individual neurons from four face patches. (**A**) Schematic diagram of experimental paradigm. Two cohorts of animals freely and repeatedly viewed the same set of naturalistic videos, during single unit recording and fMRI acquisition, respectively. The two measures of neural activity could be directly correlated, as the responses of both were synchronized to the same video content. (**B**) Schematic diagram of microwire bundle recording locations in this study, in all cases introduced into fMRI-identified of face patches. Animal identifications for each face patch recording are listed with hemispheric information in parenthesis. Abbreviation for face patches: ML, middle lateral; AF, anterior fundus; pAM, posterior part of anterior medial; aAM, anterior part of anterior medial. (**C**) Representative MR images showing recording electrode in each face patch. **(D**) Basic multimodal analyses underlying single-unit fMRI mapping. The time series corresponding to each neuron’s spiking fluctuations was correlated with that of fMRI voxels throughout the brain, after adjusting for the hemodynamic response profile (see **Materials and Methods**). Representative examples of single-unit correlation maps from three different neurons are shown on a sagittal section on the right. Panel (A) is created with BioRender.com.

To begin, we identified face patches in eleven behaving macaques using standard fMRI methods (*4, 17*). In nine animals, chronic microwire bundles were inserted into one or two of the four patches featured in the present study (**Fig. 1A**, see **Methods**). Each bundle consisted of flexible microwires that covered a small volume of tissue (< 1 mm^3^) and allowed us to record the activity of isolated neurons across multiple daily sessions (*16, 23*). We then recorded responses from 389 single neurons across four fMRI-defined face patches (**Fig. 1B – C**; n = 33 from middle lateral, ML, n = 224 from anterior fundus, AF, n = 77 from posterior portion anterior medial, pAM, n = 55 from anterior portion of anterior medial, aAM face patches; see **Methods** for distinction between pAM and aAM). For each neuron, we recorded the spiking time courses elicited during the repeated free viewing of naturalistic videos. For 254 of the recorded neurons, we independently measured basic category selectivity to briefly presented images of faces, objects, and other categories. Of these neurons, 166 (65%) were determined to be face-selective (see **Methods**).

Throughout the population, neighboring neurons in each face patch exhibited diverse response time courses to the same video content (see **Fig. S1A** for the time courses of all neurons in the study). In agreement with previous recordings from face patch AF (*16*), the time courses of neighboring neurons in all patches often diverged markedly, despite sharing some temporal features, even among the 166 neurons demonstrated to be face selective in separate testing (**Fig. S1A – B**). Importantly, neurons sampled from different face patches, typically several millimeters apart, often responded with very similar time courses (**Fig. S1A and C**). Together, these initial observations suggested that face-selective neurons exhibit a diversity of responses during natural modes of vision that are not segregated into distinct patches.

To explore this finding further, we applied the intermodal technique of single-unit fMRI mapping developed previously (*24*), which summarizes complex video-driven single-unit activity as a functional map of activity correlations across the brain (**Fig. 1D**). Each neuron’s unique pattern of positive and negative activity correlations sheds light on its participation in different functional brain networks under natural viewing conditions. Methodologically, this approach overlaps with paradigms mapping spontaneous neural activity using intermodal methods (*25–27*). However, in this case, correlations are driven by an external video stimulus, followed by extensive averaging across presentations, and experiments were carried out across the brains of different individuals. Thus, our method is designed to eliminate components of spontaneous or otherwise trial-unique activity and is distinctly different from conventional functional connectivity.

The temporal structure of the 15-minute natural video served as a shared timeline for electrophysiological and fMRI data collected in different animals. Response time courses were averaged across multiple video presentations, both for individual neurons (8 – 15 presentations) and for the fMRI responses (28 – 40 presentations). This averaging, combined with the inherent reproducibility observed in both modalities (*16, 17*), provided excellent estimations of the mean response time course for both neurons and voxels (also see **Fig. S1D** for examples). We focus here on the correlation of average spiking time courses with cortical voxels, only noting that structures outside the cerebral cortex, including the superior colliculus, pulvinar, striatum, amygdala, and cerebellum, were also engaged (see ref. (*24*)).

The correlation maps derived from each face patch were strikingly diverse, consistent with a previous application of this method to face patch AF (*24*). An example of one neuron’s map is shown in **Figure 2A**, where the fMRI coupling is depicted on a flattened cortical hemisphere. This neuron was recorded from face patch aAM and determined to be face-selective in separate testing (**Fig. 2A**, inset). This cortical correlation profile exhibited peaks that roughly matched the location of face patches, mapped in separate experiments (**Fig. 2A**, black lines, see **Methods**), suggesting the visual operation performed by this neuron can be related to modes of visual analysis that are largely restricted to other face patches.

**Fig. 2.**
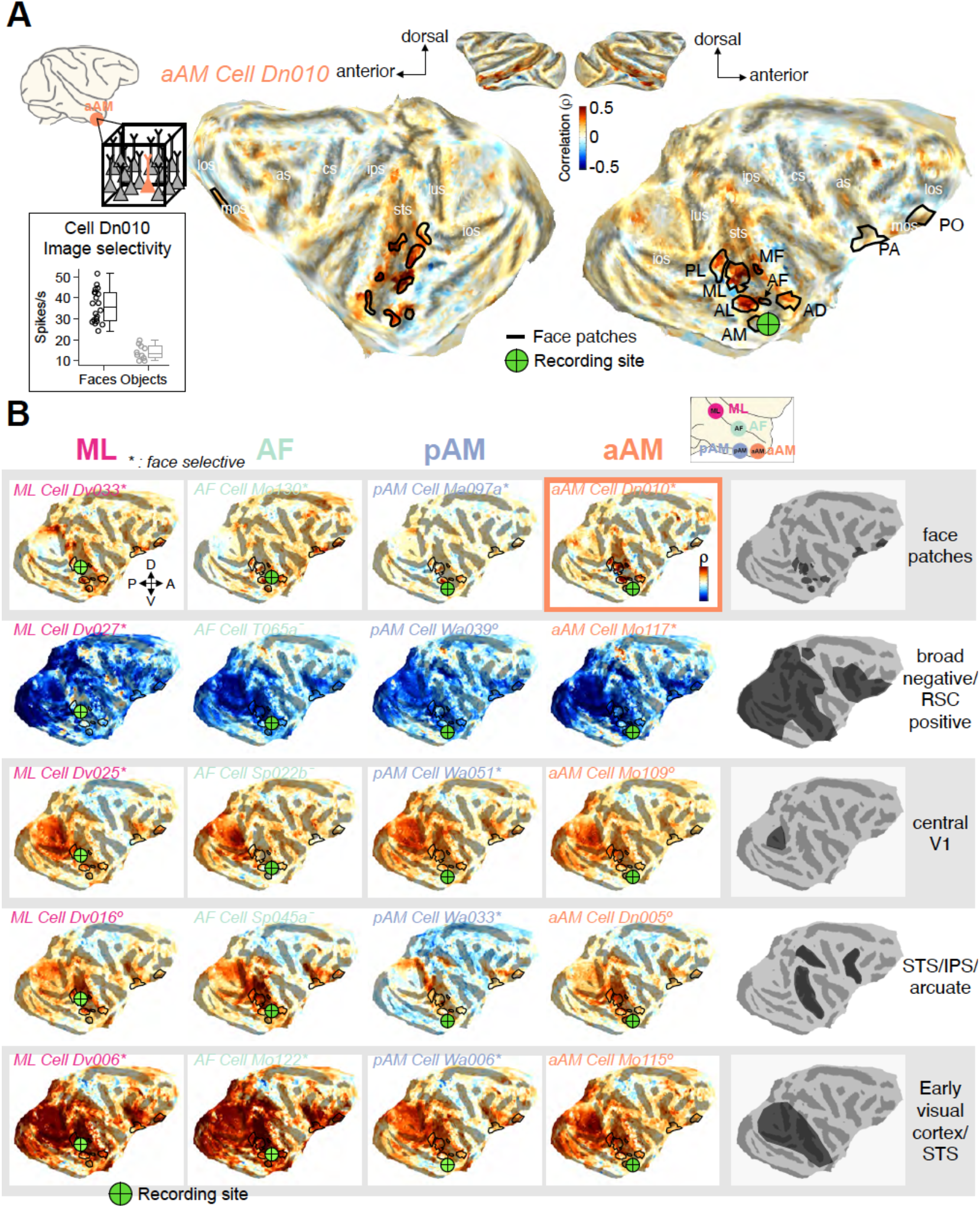
Examples of single-unit fMRI maps from neurons in four face patches. Cortical fMRI correlation pattern of example neurons recorded from four face patches projected onto the flattened cortical surface of one animal’s brain. Small green cross on each map indicates the location of recording site of the featured neuron. Location of face patches (black lines) are based on separate fMRI block design experiments. Abbreviation for face patches: PL, posterior lateral; ML, middle lateral; MF, middle fundus; AL, anterior lateral; AF, anterior fundus; AM, anterior medial; AD, anterior dorsal; PA, prefrontal arcuate; PO, prefrontal orbital face patch. (**A**) Correlation map of an example neuron from face patch aAM. Inset, spiking responses to brief presentations of faces (n = 20) and nonface objects (n = 10) demonstrate that this neuron is face selective. Center line, median; box limits, upper and lower quartiles; whiskers, range. lus, lunate sulcus; ios, inferior occipital sulcus; sts, superior temporal sulcus; ips, intraparietal sulcus; cs, central sulcus; as, arcuate sulcus; los, lateral orbital sulcus; mos, medial orbital sulcus. (**B**) Maps of 20 example neurons (cell identity is written in italics) from four face patch recording sites, presented in a matrix format. Each column of the matrix contains five neurons from each local recording site, face patches ML, AF, pAM, and aAM (from left to right), whose fMRI maps differ substantially. Each row shows fMRI maps with similar features that were common to neurons across all four recording sites, with the specific features summarized on the right. Based on separate testing, asterisks indicate neurons that were face selective, circles those that were not face selective, and dashes those that were not tested (see **Materials and Methods**). Map of the example neuron in (A) is marked with an outline. RSC, retrosplenial cortex.

The cortical correlation pattern shown in the example in **Figure 2A** was relatively common among face patch neurons, with similar examples shown from each of the four face patches in the top row of **Figure 2B**. At the same time, the columns in **Figure 2B** show that neurons with very different cortical correlation profiles were found within each of the four face patches. Dissimilar functional profiles were, for example, commonly observed among pairs of neurons recorded simultaneously from the same patch and even from the same electrode (see **Fig. S2**). Together, the rows and columns in **Figure 2B** illustrate that distinct neural subtypes were intermixed locally but shared across different face patches. For instance, each face patch contained neurons whose visual responses were broadly anticorrelated with visual areas but positively correlated with the retrosplenial cortex (**Fig. 2B**, second row), potentially suggesting that these neurons are influenced by the default mode network (*28, 29*). Each face patch also contained neurons whose activity was correlated with the foveal retinotopic cortex (**Fig. 2B**, third row), and others whose activity was correlated with dorsal stream visual areas in caudal superior temporal sulcus, intraparietal sulcus, and arcuate sulcus (**Fig. 2B**, fourth row), possibly linking to attentional network (*30, 31*). Notably absent in our analysis was any indication of elevated local fMRI correlation in the vicinity of the neural recording sites. Thus multiple, highly distinct functional profiles were shared across each of the four face patches.

We next performed a population analysis of functional profiles obtained across the four face patches. We focused on neurons determined to be face selective in separate testing (**Fig. 3**; see **Fig. S5** for the results of similar analysis applied to the entire population). To succinctly summarize the important information derived from each neuron’s correlation map, we delineated a set of 37 functional regions of interest (fROIs). For this, we identified continuous clusters of voxels across the cortical sheet that exhibited substantial correlations across all recorded neurons (see **Methods**; see **Fig. S3** for correspondence with atlas). This step allowed the correlation profile of each neuron to be expressed as a 37-element vector of value, with each element pertaining to the average correlation value within one fROI.

**Fig. 3.**
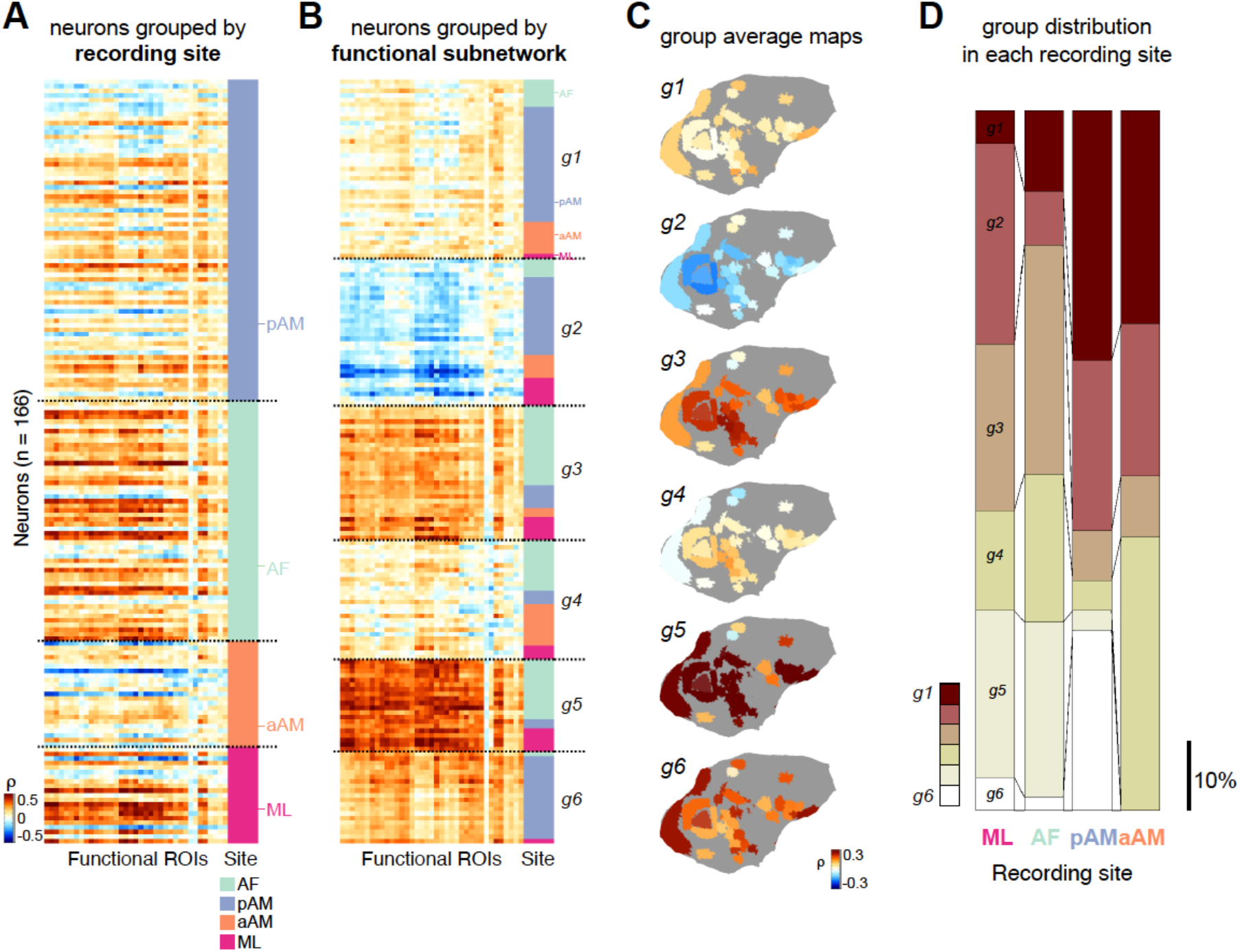
Distinct cortical correlation patterns distributed across spatially remote face patches. (**A**) fMRI correlation profiles of spiking activity from verified face-selective neurons (row; n = 166) across 37 functional regions of interest in the cerebral cortex (fROIs; see **Materials and Methods** and **Fig. S3**) (column; n = 37 from fROI 1 to fROI 37, from left to right). Each row is a 37-element vector of fROI correlation coefficients stemming from one neuron. Neurons are grouped by face patch recording location, indicated on the right in different colors. (**B**) Same format as (A), but with neurons sorted into groups *g1* – *g6* according to their functional clustering (see **Materials and Methods**). (**C**) Neural fMRI correlation profiles from (B) averaged across neurons in each group and mapped spatially onto the original fROIs on the cortical surface. (**D**) Proportion of neurons belonging to each cell group found at each of the recorded face patches. Color indicates identity of cell groups from (B) and (C). Scale bar indicates 10% of proportion.

Correlation profiles for all face-selective neurons are shown in **Figure 3A**, sorted by their recording locations. Consistent with **Figure 2B**, this figure highlights the heterogeneous composition of face cells within each patch, with marked local intermixing of neural responses to the video stimulus. To better highlight the functional groups within this matrix, we applied unsupervised (K-means) clustering to the fMRI correlation maps of the population of face-selective cells (see **Methods** and **Fig. S4**). The optimization of the clustering suggested six functional subpopulations, which we termed cell groups *g1* - *g6*. When the neural profiles were reordered based on this clustering (**Fig. 3B**), it was plain that each functional subpopulation was contributed by neurons inhabiting multiple face patches.

To better aid visualization of the functional profiles of the different cell groups, we remapped the average fROI correlation vectors from each cell group back onto the flattened brain (**Fig. 3C**). The resulting maps revealed each cell group’s specific regional coupling with the fMRI signal during naturalistic viewing. For example, one group of neurons showed relatively selective coupling to a subset of face patches, face patch AL, AD, and orbitofrontal areas (group *g1*), whereas neurons in another group (group *g4*) showed networks that appear to be complementary to the ones of group *g1*, featuring selective correlation to areas in the lower bank of STS. Another group was correlated strongly with areas in the STS and primary visual areas (group *g3*), while another (group *g6*) correlated strongly with eccentric portions of primary visual cortex in addition to the areas coupled with the group *g1*. While individual neurons composing each group differed somewhat in their specific profiles, this grouping provided an effective summary of the fMRI correlation patterns across the face-selective neural population.

Across the four patches, most functional cell groups were represented, although they were not evenly distributed (**Fig. 3D**), likely reflecting a degree of specialization of individual patches. For example, functional group *g5* with a broad, strong positive correlation profile was not observed among face-selective neurons recorded in patch aAM, nor was group *g6*. Groups *g3* and *g5*, two groups coupled strongly with STS regions, were more prominent among face-selective neurons recorded from STS face patches (ML and AF), whereas group g1 with selective face-patch correlation was observed more in face neurons from the more ventral patches (pAM and aAM).

These findings shed new light on the division of labor among single neurons within a functionally defined cortical network. They suggest that, under stimulation conditions that approximate natural vision, individual face patches are neither functionally homogeneous nor apparently arranged in a hierarchical progression. Instead, face patches overlap markedly in their functional composition. The data thus suggest the face patch system is composed of parallel threads of neural subnetworks, intermixed at a local level and carrying out distinct types of visual operations across multiple nodes of the network (**Fig. 4**).

**Fig. 4.**
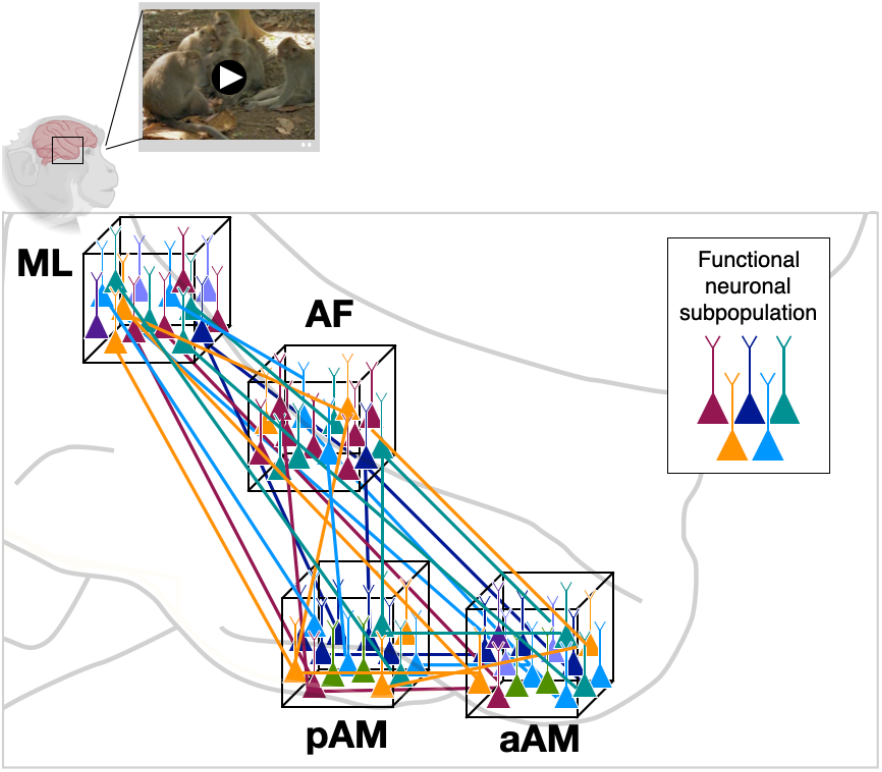
Parallel functional subnetworks pervading the face patch system. Schematic diagram depicting functionally distinct subnetworks operating in parallel within the face processing network. Local cell populations are indicated by neurons within each of the cubes, corresponding 5 to the four face patches recorded in the present study. Coloration indicates functionally distinct classes of neurons. The sharing of distinct functional operations across face patches, and the mixing of such signals locally, were central observations in the present study. The selective long-range connections among functionally similar classes of neurons (colored lines) are hypothesized.

The extreme mixing of signals in each face patch is surprising, given the common view that face patches are a clear example of domain-specificity in the brain. While this traditional view is supported by many experiments, it evidently does not extend to more natural modes of visual behavior. The difference might be attributed to the richness of the video stimuli, which include dynamic events, spatial layouts, color contrasts, and social interactions. This natural visual combination of features has seldom been incorporated into the study of face patch responses. It is also possible that active elements of vision that accompany free viewing, including exploration and attention, may unmask the different contribution of neural subpopulations (*32*). Regardless, the pronounced mixed selectivity observed during video viewing bears some resemblance to that measured in other regions across the brain (*33, 34*). It also presents practical challenges for systems and cognitive neuroscience, which often makes the tacit assumption of functional segregation in the brain.

At a more mechanistic level, it is interesting to speculate that the contrast between conventional and naturalistic testing derives from the engagement of different synaptic subpopulations providing input to individual cortical neurons. For example, during brief image presentation, a given neuron’s selectivity to faces might be conferred by the simultaneous activation of a locally shared set of inputs concerned with the analysis of facial structure. These inputs are likely concentrated within the face patches and serve as the basis for their functional definition. However, under more naturalistic conditions, the same neuron’s activity may be dominated by other, long-range connections with a sparser pattern of input among face patch neurons (see ref. (*35*) for similar ideas in different context), as well as the effects of neuromodulation during active behaviors (*36*).

Finally, combining natural conditions with novel readouts pushes researchers to explore new conceptual spaces for studying brain function. In departing from the usual practice of linking neural responses to specific visual features, less emphasis is placed on local representation and encoding and more on the physiological coordination of brain regions involved in a wide range of sensory and motor functions. The present findings underscore the value of intermodal mapping as a robust tool for characterizing neural activity during complex natural behavior, as well as the inherent value of combining controlled and naturalistic paradigms for the development of hypotheses and conceptual frameworks in systems neuroscience.

## Acknowledgments

We thank Katy Smith and Julie Hong for assistance with data collection and members in the Leopold lab for helpful comments. Functional and anatomical MRI scanning was carried out in the Neurophysiology Imaging Facility Core (NIMH, NINDS, NEI). This work utilized the computational resources of the NIH High Performance Computing Core (https://hpc.nih.gov/).

## Author contributions

Conceptualization: SHP, DAL

Data curation: SHP

Formal Analysis: SHP, KWK, BER, ENW, DBTM

Funding acquisition: DAL

Investigation: KWK, BER, ENW, DBTM

Methodology: SHP, KWK, BER, DBTM

Supervision: DAL

Visualization: SHP

Writing – original draft: SHP, DAL

Writing – review & editing: SHP, KWK, BER, ENW, DBTM, DAL

## Data and materials availability

The datasets generated and/or analyzed during the current study are available from the corresponding author on reasonable request.

## Supplementary Materials

Materials and Methods

Figs. S1 to S5

References(*37*–*47*)

## Materials and Methods

The electrophysiological and the fMRI responses analysed in the present study were recorded separately using two cohorts of animals. A full description of each set of methods, including surgical procedures, tasks, and behaviours, is given in previous publications that detail electrophysiology (*16*) and fMRI investigations (*17*) that generated part of the movie-driven datasets studied here. Technical details of the chronic microwire bundles and their targeted implantation into face patches is described in previous work (*37*). Finally, the method of single-unit fMRI mapping, comparing video-driven single unit activity and fMRI responses across the brain, is described in a previous publication (*24*).

### Subjects

Eleven rhesus monkeys (macaca mulatta, six females and five males, 5.0 – 11.3 kg, 6 – 14 years old) participated in this study. All monkeys were surgically implanted with a fiberglass headpost for head immobilization during experiments. All monkeys participated in a separate sessions of face patch localizer fMRI scans using on one or more of the following methods: a standard block design with images (*16, 17, 38*), a block design with clips of different categories (*39*), and a movie watching paradigm (*17, 38*). The locations of functionally-defined face patches had some variation across animals as often observed in previous literature (*2*), especially for anterior medial (AM) patch. In two of the animals (monkeys Dn and Mo), we found their AM patches are located approximately 2-mm anterior and lateral to the AM patches in the other animals. To differentiate these two areas, we use the term “aAM” (anterior portion of AM) to describe the anterior AM patch locations and “pAM” (posterior portion of AM) to describe the posterior AM patch locations in the other two animals (**Fig. 1B – C**). Two monkeys participated in multiple fMRI scans of free viewing of videos, whereas nine monkeys received uni- or bilateral implants of chronic microwire electrode bundles in one or two of the face patches functionally localized from the fMRI localizer scans: middle lateral (ML), anterior fundus (AF), pAM, and aAM face patches (**Fig. 1B – C**). All procedures were approved by the Animal Care and Use Committee of the US National Institutes of Health (National Institute of Mental Health) and followed US National Institutes of Health guidelines.

### Experimental Design

#### fMRI: free viewing of videos

During the functional echo planar imaging (EPI) scans, subjects viewed the visual stimuli projected onto a screen above their head through a mirror inside of the MRI scanner. Three 5-min videos depicting macaque monkeys interacting with conspecifics and humans were presented in 640 × 480 pixel resolution with a frame rate of 30 frames per second. Each session started with an eye calibration and the calibration was repeated every two 5-minute trials. Each video viewing began with a 500-ms presentation of a white central point surrounded by a 10° diameter annulus indicating the gaze-position tolerance window, which was approximately the size of the video stimulus. Subjects received a drop of juice reward every 2 s as long as their eyes were within the movie frame.

#### Electrophysiology: free viewing of videos

Nine electrophysiology subjects viewed the same three 5-min videos used in fMRI video watching experiments during the electrophysiological recordings. The videos were presented on an LCD or OLED monitor within a rectangular frame of 10.4° wide and 7.6° high. Eye position was calibrated at the beginning of each session and as needed during subsequent inter-trial intervals. One monkey viewed the movie without liquid reward, whereas the other eight received a water or juice reward for directing gaze to the movie, defined as a large “fixation window” of 10.5°.

#### Electrophysiology: fixation on flashing images

Six of nine electrophysiology subjects underwent another set of passive fixation experiment with images of different categories on each day before their video watching experiments. Subjects viewed a stimulus set consisting of 60 images containing sets of face categories (monkey faces and human faces) and non-face categories (objects and scenes). Images were 12° in width and randomly presented for 100 ms with 200-ms blanks. Subjects were rewarded every six presentations of images for maintaining their gaze within 1.2° – 2° in radius of the fixation point (size: 0.2° radius).

### Data Acquisition

#### Whole-brain fMRI

Structural and functional images were acquired in a 4.7-T, 60-cm vertical scanner (Bruker Biospec, Ettlingen, Germany) equipped with a Bruker S380 gradient coil in the Neurophysiology Imaging Facility Core (NIMH, NINDS, NEI). From two female monkeys, we collected whole brain images with an eight-channel transmit and receive radiofrequency coil system (Rapid MR International, Columbus, OH). Functional EPI scans were collected as 40 sagittal slices covering the whole brain with an in-plane resolution of 1.5 mm × 1.5 mm and a slice thickness of 1.5 mm. Repetition time and echo time were 2.4 s and 12 ms, respectively. Monocrystalline iron oxide nanoparticles (MION), a T2* contrast agent, was administered prior to the start of EPI data acquisition. Subjects participated in multiple fMRI scans (28 – 40) of free viewing of three 5-minute videos across multiple days. All aspects of the task related to timing of stimulus presentation, eye-position monitoring, and reward delivery were controlled by custom software courtesy of David Sheinberg (Brown University, Providence, RI) running on a QNX computer. Horizontal and vertical eye positions were recorded using an MR-compatible infrared camera (MRC Systems, Heidelberg, Germany) fed into an eye tracking system (SensoMotoric Instruments GmbH, Teltow, Germany) at the sampling rate of 200 Hz.

#### Electrophysiological recordings from cortical face patches

From nine monkeys, electrophysiological recordings were obtained from bundles of 32, 64, or 128 NiCr microwires chronically implanted in one or more face patches. The microwire electrodes were designed and initially constructed by Dr. Igor Bondar (Institute of Higher Nervous Activity and Neurophysiology, Moscow, Russia) and subsequently manufactured commercially (Microprobes). Data were collected using either a Multichannel Acquisition Processor with 32-channel capacity (Plexon) or a neural recording system with 128-channel capacity (Tucker-Davis Technologies). All aspects of the task related to timing of stimulus presentation, eye-position monitoring, and reward delivery were controlled either by a custom software running on a QNX computer (courtesy of David Sheinberg (Brown University, Providence, RI)) or by NIMH MonkeyLogic(*40*). Horizontal and vertical eye positions were recorded using an infrared video-tracking system (EyeLink; SR Research). Event codes, eye positions and a photodiode signal were stored to a hard disk using OpenEX software (Tucker Davis Technologies).

### Data Analysis

#### Pre-processing of fMRI data

All fMRI data were analysed using the AFNI/SUMA software package (*41*) (https://afni.nimh.nih.gov/afni) and custom-written MATLAB code (MathWorks, Natick, MA). Raw images were first converted into AFNI data file format. Each scan underwent slice-timing correction, correction for static magnetic field inhomogeneities using the PLACE algorithm (*42*), motion correction (AFNI function 3dvolreg), and between-session spatial registration to a single EPI reference scan. The first seven time frames of each scan were discarded for T1 stabilization, followed by removal of low-order temporal trends (polynomials up to the 5th order). The time series of each voxel was converted to units of percent signal change by subtracting, and then dividing by, the mean intensity at each voxel over the course of each 5-minute scan. The high-resolution anatomy volume was skull stripped and normalized using AFNI and then imported into CARET (*43*). Within CARET, surface and flattened maps were generated from a white matter mask and exported to SUMA for viewing.

#### Pre-processing of electrophysiology data

Individual spikes were extracted off-line from the broadband signal (2.5-8 kHz) after band-pass filtering between 300 and 5000 Hz for spikes, using the OfflineSorter software package (Plexon) or WaveClus spike-sorting package (*44*). We identified single units longitudinally across 2 – 5 days based on multiple criteria, including the similarity in waveform shapes and inter-spike interval histograms across days, and the distinctive visual response pattern generated by isolated spikes as a neural “fingerprint,” which we previously found in this area of cortex. Further details on spike sorting and longitudinal identification of neurons across days can be found in ref. (*23, 37*). In total, we isolated 389 single units from four face patches: n = 33 from the patch ML (monkey Dv), n = 224 from AF (48 from monkey To, 5 from Ro, 16 from Si, 106 from Sp, 49 from Mo), n = 77 from pAM (27 from monkey Ma, 50 from Wa) and n = 55 from aAM (13 from monkey Dn, 42 from Mo).

#### Computation of single-unit functional maps of individual neurons

For each of the 389 neurons recorded from four face patches, we computed whole-brain functional maps, where the value of each voxel was the correlation coefficient between its fMRI time course and the single neuron’s time course (**Fig. 1D**). The detailed methods are described in a previous publication (*24*). Briefly, for single-unit responses, we down-sampled each single-unit response to match the fMRI temporal resolution (2.4 second), averaged time courses across trials for a given 5-minute movie, convolved a generic hemodynamic impulse response function to the averaged and normalized (i.e. centered to the mean) time course, then concatenated the convolved neuronal time courses from the three different movies, resulting in a single 15-minute time series for each neuron. For fMRI responses, we averaged the time series across all trials for each 5-minute movie, then concatenated across three movies, resulting in 15-min (i.e. 375 volumes) time series of each voxel in the whole brain. After this preparation of the time courses, for each single unit we computed the Spearman’s rank correlation coefficients between the neuronal time course and fMRI time courses of all the voxels (n = 27,113) in the whole brain.

#### Principal component analysis on neuronal time series

To assess the diverse nature of video-driven responses of the population of neurons, we applied the principal component analysis on neuronal time series of 15-minute. Specifically, for each neuron we computed the number of spikes in a non-overlapping time bin of 2.4 second from single trial for a 5-minute movie (i.e. 125 data points), then averaged the time series across trials. Then, we concatenated the averaged time course across three movies, generating a single 15-minute time series for each neuron (i.e. 375 data points). Each neuron’s time series was then converted into z-score by subtracting its mean and dividing by standard deviation. The matrix of the normalized time series from the whole population (neuron × time) was then fed into the pca function of MATLAB (pca.m), where the Singular Value Decomposition was applied. The projection value on the first principal component axis was used to sort the neurons along the first principal component as shown in **Fig. S1A**.

#### Computation of face selectivity

For 254 neurons (n = 33 from ML, n = 89 from AF, n = 77 from pAM, and n = 55 from aAM) recorded from six of the nine electrophysiology subjects that underwent passive fixation experiments on different categories of images, we evaluated the categorical face selectivity (*3, 45–47*). First, responses to each image exemplar were calculated as the average spike rate in a fixed window of 0 – 200 ms after stimulus onset, and baseline subtracted using the average firing rate in a window of -50 – 0 ms from stimulus onset across all image exemplars. Then, we computed a face selectivity index (FSI) from the mean responses to faces and nonface objects following the conventions of previous studies (*46, 47*) as below:

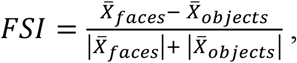

where FSI varies between -1 and 1, with additional condition of FSI = -FSI when 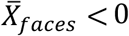 and 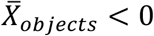 to incorporate inhibitory face-selective response. When the FSI of a neuron is larger than 0.33 (i.e. twice larger excitatory or inhibitory responses to the images of faces than the images of nonface objects), the neuron is defined as “face-selective” (e.g. neurons marked with black in **Fig. S1A** and **Fig. S5D**, example neurons marked with asterisks in **Fig. 2B** and **Fig. S2**, and neurons included for **Fig. 3**). In total, 166 neurons out of 254 neurons (65%) were determined face selective (n = 21 from ML, n = 52 from AF, n = 70 from pAM, n = 23 from aAM).

#### Clustering of single units based on the whole-brain correlation maps

To group the neurons based on their whole-brain movie-driven correlation pattern irrespective of their recording location, we applied a k-means clustering algorithm on the matrix of correlation values between the face-selective neurons (n = 166) and within-brain voxels (n = 27,113). We used the kmeans.m function in MATLAB’s Statistics Toolbox with the squared Euclidean distance metric. We repeated the clustering procedure 100 times each for K values ranging from 2 to 40, and for each repetition for each K, we computed the percentage of variance explained by the clustering (i.e., the between-clusters sum-of-squares relative to the total sum-of-squares) (**Fig. S4A**). As the K value increased, there was an asymptotic increase in explained variance as expected, but no clear point of concavity. By computing the amount of increase in explained variance we gain as we increase K (i.e. difference in explained variance between K-1 and K, **Fig. S4B**), we found that there was much less gain after increasing K from 5 to 6 (arrow in **Fig. S4B**). Since K-means clustering is not a deterministic algorithm but a stochastic one, we evaluated the stability of the clustering by computing the standard deviation of explained variance across the 100 repetitions (**Fig. S4C**). The variability of the explained variance showed an asymptotic increase as K increases as expected. However, for the case of K = 6 it started to become more stable (i.e. less variable) and showed the most stability comparing to the cases of Ks around it. Based on these two additional factors (**Fig. S4**), we determined that six clusters, or groups of neurons, were appropriate to summarize this population of neurons. For the entire population of neurons (n = 389), we performed identical procedure on clustering and selection of the number of clusters (**Fig. S5A – C**), resulted in determining ten clusters were appropriate to summarize the entire population of all the neurons irrespective of their selectivity for faces (**Fig. S5D – E**).

#### Delineation of functional regions of interest (fROIs) on cortex

To summarize the correlation profile of each neuron with cortical regions, we delineated a set of 37 functional regions of interest (fROIs) on the cortex of one fMRI animal (Fig. S3). For a broad and fair inclusion of cortical regions that are correlated with face patch neurons regardless of their recording sites, we first computed which cortical regions are correlated with face patch neurons higher than the chance level (24). More specifically, for each voxel, the fraction of neurons that exhibit an absolute correlation value higher than 0.3 is computed and mapped as shown in the Fig. S3A. The population map of fraction of highly correlated neurons, where the fraction is computed using the total number of neurons (n = 389), provided a useful and complementary guidance on which cortical regions are correlated with the cells we recorded. Because the fraction of neuron map has no bearing on which voxels are correlated together (i.e. specific patterns of correlation or which voxels can be grouped together as one fROI), we additionally considered the repeated patterns of correlation observed across individual neurons. We first used SUMA to draw the ROIs on the cortical surface (Fig. S3B) and then used AFNI for a necessary correction of the voxels for each ROI in 3D slices (Fig. S3C). The resulting 37 fROIs covered the occipital, temporal, parietal, and frontal lobes. Most of the fROIs overlap with multiple cytoarchitectonic areas, the correspondence of which has been established by a separate registration of this fMRI animal’s brain to D99 atlas using AFNI (see Fig. S3C for correspondence with atlas), as our main purpose of the ROI delineation was an effective summary or description of each neuron’s cortical correlation profiles.

**Fig. S1.**
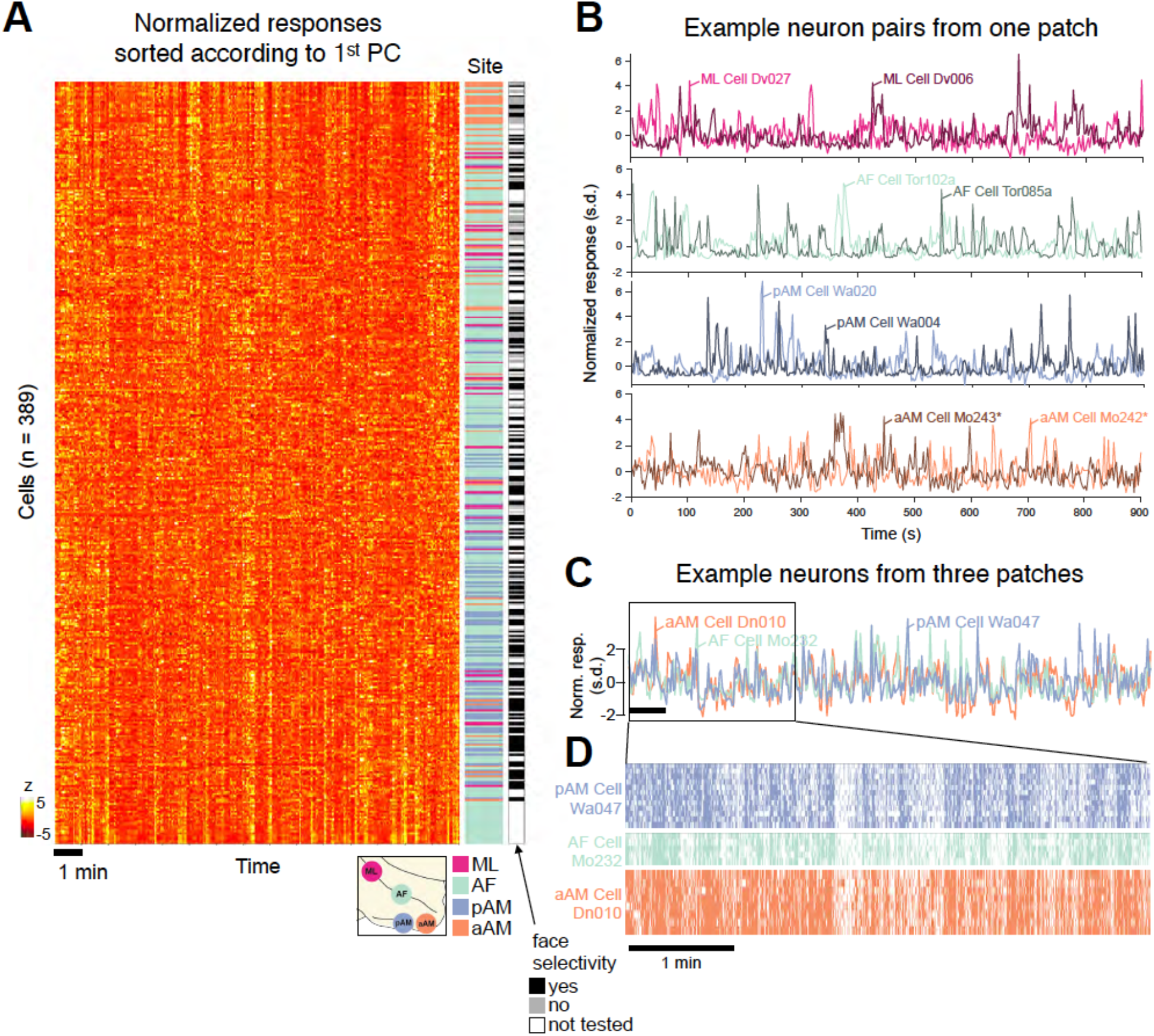
Movie-driven response time course of neurons from four face patches. (A) Left: a heat-map depicting normalized movie-driven response time course of all the neurons (rows), sorted according to the first principal component (see **Materials and Methods**). Scale bar: 1 minute. Right: electrode location (color) and face selectivity (grayscale) are indicated for each neuron. (**B**) Time courses of four example pairs of neurons recorded from one patch (from ML, AF, pAM, and aAM from top to bottom). Asterisks on the last pair indicate that this pair of neurons are recorded from the same electrode. (**C**) Time courses of three example neurons recorded from three face patches showing shared temporal structure. Normalized (z-scored) responses of each neuron are shown. Each time course is an average across multiple viewings of the same video. (**D**) Raster plots of each of three neurons shown in c across multiple viewings of a five-minute video. Scale bars in (C) and (D) are 1 minute.

**Fig. S2.**
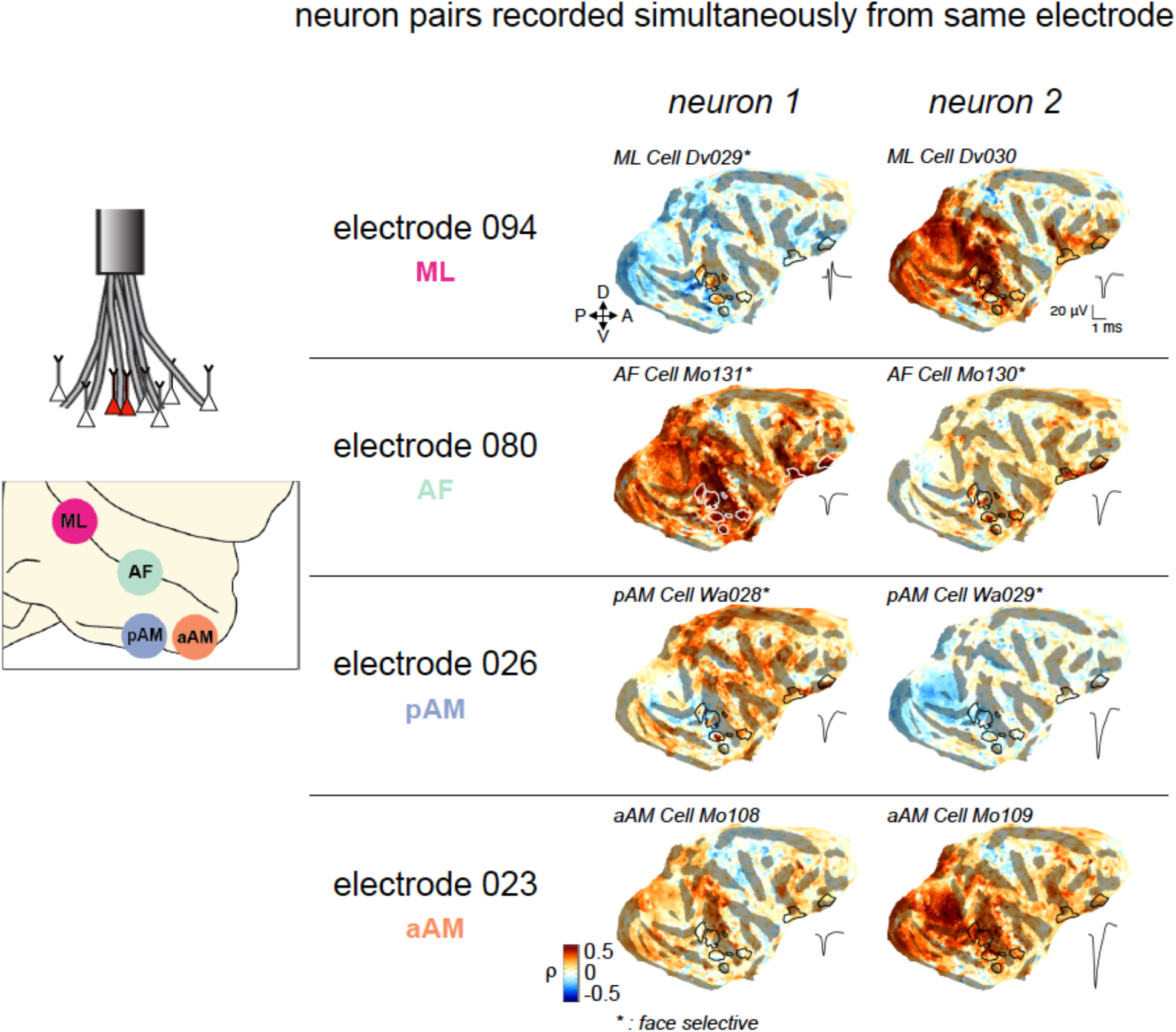
Whole-brain correlation maps of four example pairs of neurons recorded simultaneously from same electrode. Each pair is taken from one of the four patches. Waveform of each single unit is shown on the right side of the correlation map. Based on separate testing, asterisks indicate neurons that were face selective. Scale bar for waveforms: 20uV, 1 ms.

**Fig. S3.**
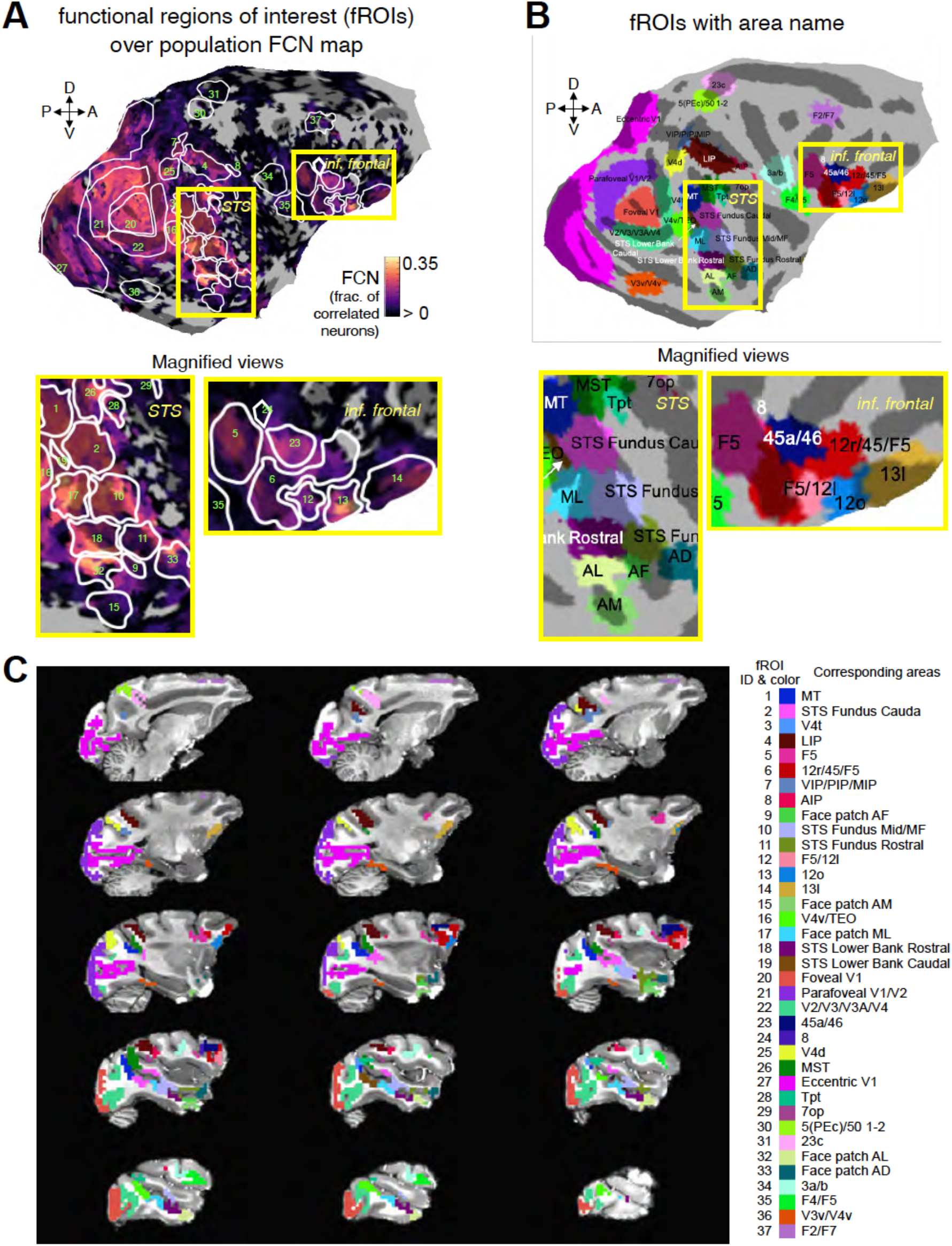
Cortical functional regions of interest (fROIs). (**A**) fROIs superimposed on the fraction of correlated neuron (FCN) map. FCN was computed as the proportion of neurons that exhibit absolute correlation coefficient higher than 0.3 (see **Materials and Methods**) Boundaries of 37 fROIs with identification number is shown on top of the population FCN map. Voxels without color indicate that none of the 389 neurons recorded show absolute correlation higher than 0.3 with them. Identification number is identical as the fROIs are ordered in the correlation matrix shown in **Fig. 3**. Regions around superior temporal sulcus (STS) and inferior frontal part (inf. frontal) are magnified at the bottom. (**B**) fROIs on the flattened cortical surface of one fMRI animal. 37 fROIs are visualized with different colors, with names of corresponding areas for each fROI. (**C**) Left, fROIs are visualized on sagittal slices of structural MR images. Right, identification numbers (same as (B)) and color (same as a) of fROIs are shown with names of corresponding areas from D99 atlas.

**Fig. S4.**
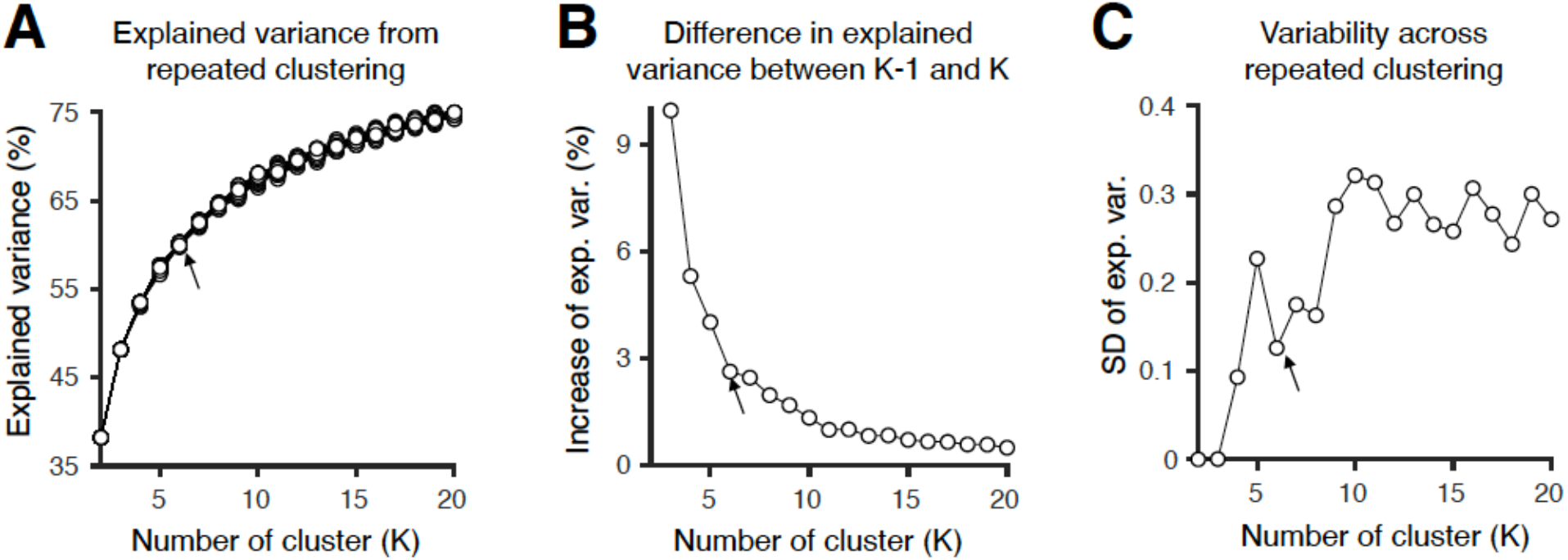
Unsupervised (K-means) clustering of single cells based on whole-brain correlation profiles. (**A**) Explained variance as a function of the number of clusters (K). Individual lines and markers are from each clustering event that is repeated for 100 times (see **Materials and Method**s). The case of K = 6 is marked with arrow. (**B**) Difference in explained variance between K-1 and K. The difference between the explained variance in the case of K = 5 and in the case of K = 6 is marked with arrow. (**C**) Variability across repeated clustering. Standard deviation (SD) of explained variance across 100 clustering occurrences for a given K is shown as a function of K. The case of K = 6 is marked with arrow.

**Fig. S5.**
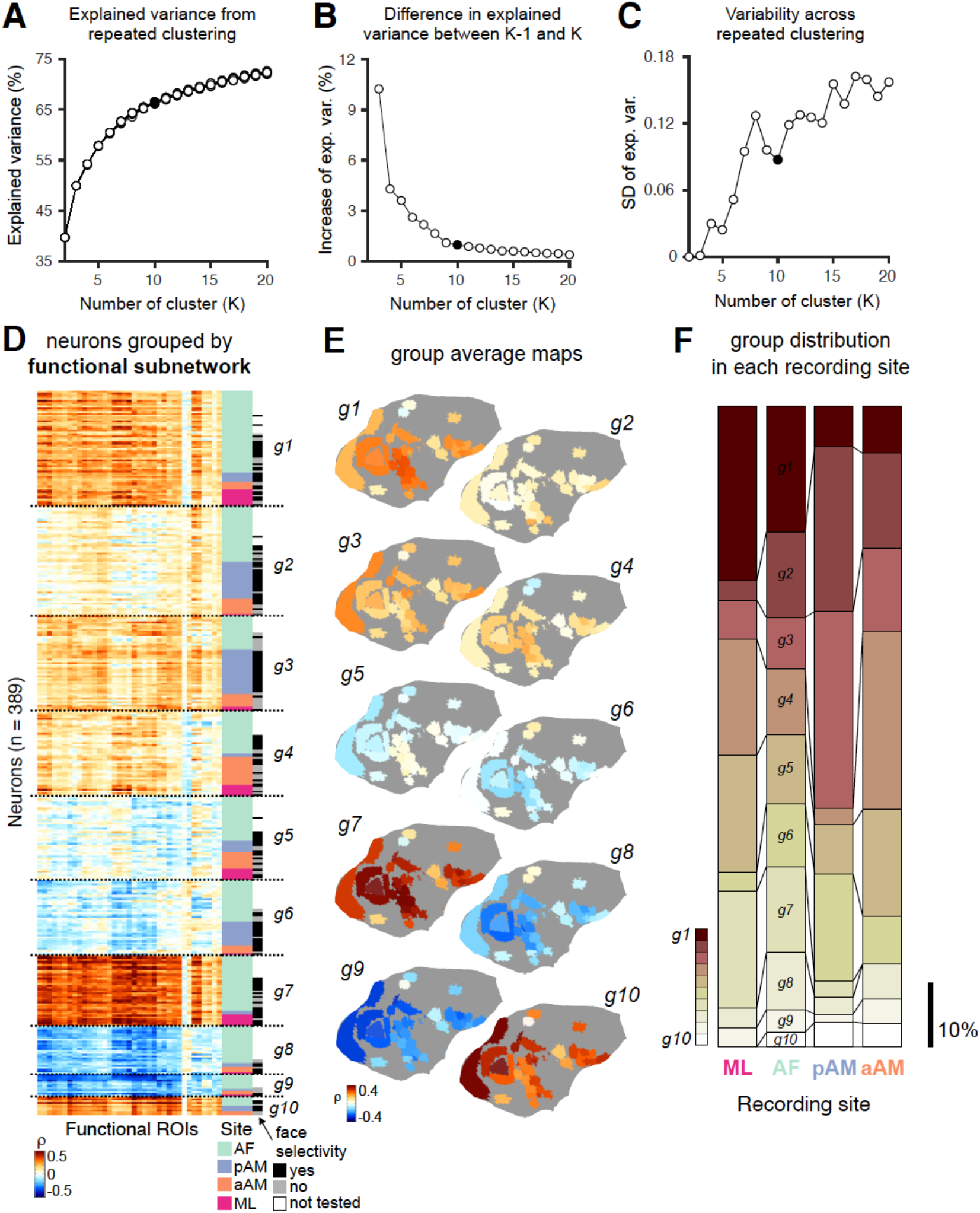
Unsupervised clustering of all the neurons (n = 389) from four face patches based on whole-brain correlation profiles. (**A – C**) Evaluation of repeated clustering performance and stability as a function of the number of cluster (K). Same format as **Fig. S4**. The case of K = 10 is marked as black. (**D – F)** Depiction of distinct ten functional groups of neurons (*g1* – *g10*) in correlation matrix with indications of recording site (color) and face selectivity (grayscale) from separate testing for each neuron (**D**), average group maps (**E**), and their distribution in each recording site (**F**). Same format as **Fig. 3B – D**.

